# Variation of HbA1c affects cognition and white matter microstructure in healthy, young adults

**DOI:** 10.1101/507186

**Authors:** Jonathan Repple, Greta Karliczek, Susanne Meinert, Katharina Förster, Dominik Grotegerd, Janik Goltermann, Volker Arolt, Bernhard Baune, Udo Dannlowski, Nils Opel

## Abstract

The metabolic serum marker HbA1c has been associated with both impaired cognitive performance and altered white matter integrity in patients suffering from diabetes mellitus. However, it remains unclear if higher levels of HbA1c might also affect brain structure and function in healthy subjects. With the present study we therefore aimed to investigate the relationship between HbA1c levels and cognitive performance as well as white matter microstructure in healthy, young adults.

To address this question, associations between HbA1c and cognitive measures (NIH Cognition Toolbox) as well as DTI-derived imaging measures of white matter microstructure were investigated in a publicly available sample of healthy, young adults as part of the Humane Connectome Project (n= 1206, mean age= 28.8 years, 45.5 % male).

We found that HbA1c levels (range 4.1-6.3%) were significantly inversely correlated with measures of cognitive performance. Higher HbA1c levels were associated with significant and widespread reductions in fractional anisotropy (FA) controlling for age, sex, body-mass index and education. FA reductions were furthermore found to covary with measures of cognitive performance. The same pattern of results could be observed in analyses restricted to participants with HBAlc levels below 5.7%.

The present study demonstrates that low-grade HbA1c variation below diagnostic threshold for diabetes is related to both cognitive performance and white matter integrity in healthy, young adults. These findings highlight the detrimental impact of metabolic risk factors on brain physiology and underscore the importance of intensified preventive measures independent of the currently applied diagnostic HbA1c cut-off scores.

**Highlights:**

- HbAlc levels are negatively associated with cognitive performance in healthy, young adults in a sample of n=1206 (Human Connectome Project).
- HbA1c levels even below strict prediabetes thresholds are negatively associated with white matter integrity throughout the brain.
- White matter integrity in affected regions is positively associated with cognitive performance.

## 1. Introduction

With an estimated global prevalence of 8.5% diabetes mellitus represents one of the most significant health care issues of modern times. Diabetes is one of the leading causes for several highly prevalent cardiometabolic diseases including myocardial infarction, kidney failure and ischemic stroke ^1^. HbA1c, which represents the percentage of glycated Hemoglobin and is thought to resemble the three-month average plasma glucose concentration, is considered as the gold standard for the control of diabetes treatment efficacy. Recently, its role in aiding primary type 2 diabetes diagnoses has been emphasized by the World Health organization. A HbA1c percentage of 6.5% was considered a reliable cutoff for the diagnosis of diabetes, with a high-risk range of 6-6.5% ^2^. The American Diabetes Association proposes even stricter levels and suggests that HbA1c levels above 5.7% should be considered as a biomarker for prediabetes^3^.

While diabetes affects physiological function of multiple organ systems, its devastating impact on brain physiology has only recently been further elucidated. Besides ischemic stroke, diabetes and impaired glucose metabolism increases the risk for both vascular and Alzheimer’s dementia ^4,5^. Interestingly, there is also robust evidence for a negative association between HbA1c and both cognitive function as well as brain structural integrity in patients with type 2 diabetes without dementia ^6^. This finding is further supported by recent research suggesting accelerated deterioration of measures of processing speed in type 1 diabetes patients and its association with glycemic control in these patients^7^. Moreover, results from young type 1 diabetes patients show, that lower HbA1c is associated with higher fractional anisotropy (FA), a widely employed neuroimaging marker of white matter microstructural integrity derived from Diffusion Tensor Imaging (DTI)^8,9^. Similarly, a relationship of increased HbA1c and decreased fractional anisotropy, indicating white matter damage, as well as reduced neurocognitive performance has been shown in type 1 diabetes patients ^7,10^. This notion of a relationship between altered white matter integrity and cognition is supported by further reports ^11,12^, with most robust evidence for the cognitive subdomains of processing speed ^13,14^ and executive function ^15,16^.

However, to the best of our knowledge, associations of HbA1c, cognition and white matter integrity have not been examined in healthy subjects so far. More precisely, it is unknown if variation of HbA1c levels even below established diagnostic cut-off scores associate with both cognitive performance and white matter microstructure. Considering that individual glucose metabolism and HbA1c levels in general are modifiable risk factors, solving this research question would have important consequences for future preventive efforts.

To address this currently unresolved research question we aimed to investigate potential associations between HbA1c, cognitive performance and white matter microstructure in a well powered sample of healthy, young adults. We hypothesized a) that HbA1c would be associated with reduced cognitive performance, b) that HbA1c would be associated with reduced FA and c) that cognitive performance would correlate negatively with FA. Furthermore, as both the prevalence of metabolic syndrome ^17^ as well as the level of cognitive performance ^18^ are strongly associated with socioeconomic status, we investigated whether the aforementioned hypothesized associations persisted when correcting for total years of education as a proxy for socioeconomic status^19^.

## 2. Material and Methods

### 2.1 Participants

We investigated open-access brain imaging data from the Human Connectome Project (HCP) WU-Minn HCP 1200 Subjects Data Release ^20^ (for further information on details of data acquisition and processing in this sample please see: https://www.humanconnectome.org/study/hcp-young-adult/document/1200-subjects-data-release). Exclusion criteria were neurodevelopmental disorders (e.g., autism), documented neuropsychiatric disorders (e.g., schizophrenia or depression), neurologic disorders (e.g., Parkinson’s disease), diabetes or high blood pressure ^20^. All 1206 subjects were included in the present study. Analyses were performed with the maximum number of available data for each analysis. We report the respective available n for each analysis. Five subjects were excluded, as their HbA1c levels were outside 3 standard deviations of the mean value.

The mean age of the sample was 28.8 years (standard deviation (SD)= 3.69), 45.5 % of all participants were male. Body Mass Index (BMI) was calculated as the body weight (in kilogram) divided by squared body height (in meters) (mass_kg_/height_m_^2^). BMI was available for n= 1200, the mean was 27.09 (SD= 5.89). Total education years, a proxy for socioeconomic status, were calculated as the years of completed education (range 11-17 years, mean= 14.87 years). Subjects were primarily recruited in Missouri; additional recruiting efforts were made to ensure that participants generally reflect the ethnic and racial composition of the U.S. population. For data acquisition participants visited the Washington University, St. Louis, Missouri, twice with a fixed order of MRI scanning and extensive behavioral assessment on each day^20^.

### 2.2 HbA1c analysis

To determine HbA1c levels, blood samples were taken into EDTA test tubes and the turbidimetric inhibition immunoassay for hemolyzed whole blood was applied. After adding the buffer/antibody reagent, the glycated hemoglobin A1c reacts with anti-HbA1c antibodies forming soluble antigen-antibody complexes. Subsequently, the buffer/polyhapten reagent is added, so polyhaptens and excess anti-HbA1c antibodies react to an insoluble complex, which is measured turbidimetrically. Hemoglobin is quantified bichromatically in form of a derivative with a characteristic absorption spectrum. Finally, the HbA1c percentage is calculated as follows: HbA1c [%] = (Raw HbA1c Mass/Raw Hemoglobin) x 91.5 + 2.15 ^21^. HbA1c was available for n= 820, the mean was 5.24% (range 4.1-6.3%). 791 subjects had HbA1c scores of 5.7% or below.

### 2.3 Cognitive performance

Cognitive measures were available for n= 1187 subjects. All cognition tests included in the HCP dataset were included in the study. The NIH Cognition Total Composite Score (“global cognition score”) is calculated as the average of subtest-scores contained in the NIH Toolbox Cognition Battery (Flanker, Dimensional Change Card Sort, Picture Sequence Memory, List Sorting and Pattern Comparison, Picture Vocabulary and Reading Recognitions) and reflects overall cognitive performance ^18,22,23^. Scores underwent age-adjusted normalization, so that a score of 100 indicates an average performance within the age group and a plus or minus of 15 represent 1 SD above or below. For further information on all cognitive measures see Supplementary Methods 1.

### 2.4 DTI data acquisition

DTI data was available for n=1050 in the HCP sample. Data for the HCP was acquired on a customized Siemens 3T “Connectome Skyra” housed at Washington University in St. Louis, using a standard 32-channel Siemens receive head coil and a “body” transmission coil designed by Siemens specifically for the smaller space available using the special gradients of the WU-Minn and MGH-UCLA Connectome scanners ^21,24^.

A full diffusion MRI session includes 6 runs (each approximately 9 minutes and 50 seconds), representing 3 different gradient tables, with each table acquired once with right-to-left and left-to-right phase encoding polarities, respectively. Each gradient table includes approximately 90 diffusion weighting directions plus 6 b=0 acquisitions interspersed throughout each run. Diffusion weighting consisted of 3 shells of b=1000, 2000, and 3000 s/mm2 interspersed with an approximately equal number of acquisitions on each shell within each run (Sequence: Spin-echo EPI, TR 5520 ms, TE 89,5 ms, flip angle 78 deg, refocusing flip angle 160 deg, FOV 210×180 (RO × PE), matrix 168×144 (RO × PE), slice thickness 1.255 mm, 111 slices, 1.25 mm isotropic voxels, multiband factor 3, echo spacing 0.78 ms, BW 1488 Hz/Px, phase partial fourier 6/8, b-values 1000, 2000, 3000 s/mm2) ^24,25^.

### 2.5 DTI data preprocessing and analysis

Diffusion data accessible from the HCP were preprocessed with their MR Diffusion Pipeline ^26^, which normalizes the b0 image intensity across runs; removes EPI distortions, eddy-current-induced distortions, and subject motion; corrects for gradient-nonlinearities; registers the diffusion data with the structural; brings it into 1.25mm structural space; and masks the data with the final brain mask: 1. Basic preprocessing: Intensity normalization across runs, preparation for later modules. 2. ‘TOPUP’ algorithm for EPI distortion correction. 3. ‘EDDY’ algorithm for eddy current and motion correction. 4. Gradient nonlinearity correction, calculation of gradient bvalue/bvector deviation. 5. Registration of mean b0 to native volume T1w with FLIRT BBR+bbregister and transformation of diffusion data, gradient deviation, and gradient directions to 1.25mm structural space. The brain mask is based on FreeSurfer segmentation.

Tract-based spatial statistics (TBSS)^27^ is a well-established analysis method for DTI imaging and was applied as described extensively in a recent study^28^. Briefly, standard TBSS preprocessing was performed ^27^: The FA images were registered to the FMRIB58 FA template and averaged to create a mean FA image. A WM skeleton was created with an FA threshold of 0.2 and overlaid onto each subject’s registered FA image. Individual FA values were warped onto this mean skeleton mask by searching perpendicular from the skeleton for maximum FA values.

To test for statistical significance, we used the non-parametric permutation testing implemented in FSL’s ‘randomize’ with 5000 permutations. Threshold-Free Cluster Enhancement (TFCE) ^29^ was used to correct for multiple comparisons. This allows to estimate cluster sizes corrected for the family-wise error (FWE; p < .05, 5000 permutations). MNI coordinates and cluster size at peak voxel were derived with FSL Cluster and the corresponding WM tract retrieved from the ICBM-DTI-81 white-matter atlas^30^.

### 2.6 Statistical analyses

1. First, to investigate potential associations between HbA1c and cognitive measures, we performed correlational analyses in SPSS (IBM Version 25):

a. As the primary analysis, the correlation between HbA1c and the global cognition score was assessed. Furthermore, additional correlational analyses between HbA1c and all available measures of cognitive subdomains were carried out to further delineate the contribution of specific subdomains to the observations made in step a) above. For more details on these tests please see Supplementary Material 1. To further test for specificity of HbA1c related effects, analyses were repeated by including BMI as nuisance variable in a partial correlation. To rule out that our findings were mainly driven by observations above the diagnostic threshold for diabetes, the analysis was repeated by excluding all subjects with HbA1c scores higher than 5.7%.
2. Second, we tested for linear effects of HbA1c on FA across the entire WM skeleton with general linear models within FSL. We accounted for the effects of nuisance covariates that could influence WM structure: age ^31^ and sex ^32^. Again, analyses were repeated by including BMI as additional nuisance variable and by excluding all subjects with HbA1c scores higher than 5.7%.
3. Third, we sought to investigate the relationship between cognition and HbA1c-related white matter microstructure. In line with analysis step 1 above, associations between FA and the global cognition score were assessed in a first step. Additionally, correlational analyses between FA and measures of cognitive subdomains were carried out:

a. Linear effects of total cognitive performance on FA across the WM skeleton were analyzed with a general linear model. Following our hypothesis focusing on HbA1c related FA, a mask of all significant voxels from the HbA1c ~ FA analyses (2.) was applied for this analyses step. Age and sex were included as nuisance variables.
b. We extracted one FA value, which was the mean of all significant voxels in this analysis. With this FA value, we performed exploratory correlational analyses with all measures of cognitive subdomains available in the HCP data set. For more details on these tests please see Supplementary Material 1.
4. Finally, we aimed to assess the influence of education as a proxy for socioeconomic status on results of the aforementioned analyses. To this end, we first investigated the direct association of total education years on HbA1c, cognition and extracted FA values and then repeated the analyses performed in 1., 2. and 3. while additionally correcting for years of education.

## 3 Results

### 3.1. HbA1c and cognitive performance

HbA1c scores were significantly negatively correlated with the global cognition score (n= 806, r= -.108, p= .002). Likewise, significant inverse correlations between HbA1c levels and several cognitive subdomains could be observed with strongest effect sizes for the List Sorting Working Memory test (working memory) (df= 730, r= -.113, p= .002), the Picture Vocabulary test (vocabulary knowledge) (df= 730, r= -.125, p= .001) and the Penn Progressive Matrices test (fluid intelligence) (df= 803, r= -.118, p= .001). For an overview of the respective analyses of all cognitive subdomains, please see Table 1. The correlation between HbA1c and the global cognition score remained significant when correcting for BMI (df= 774, r= -.074, p= .036), but was reduced to trend-level significance in an analysis restricted to subjects with an HbA1c<= 5.7 (n= 777, r= -.067, p= .064) while still significant associations could be observed for the Picture Vocabulary test in these analyses (Picture Vocabulary test: n= 791, r= -.084, p= .018; Penn Progressive Matrices test: n= 788, r= -.064, p= .074; List Sorting Working Memory test: n= 791, r= -.041, p= .255).

### 3.2. HbA1c and white matter microstructure

Complete HbA1c and DTI data were available for a sample of n= 737 subjects. We found a significant negative correlation (p_FWE_-corrected< .05) of FA and HbA1c, correcting for age and sex, in large, widespread clusters, including the genu of the corpus callosum, the bilateral longitudinal superior fascicle, the bilateral internal and external capsule, the bilateral uncinate fascicle, the corticospinal tract and the cerebellar peduncles among others (Supplementary Results 1, Figure 1).

This negative correlation remained significant when further correcting for BMI (p_FWE_-corrected< .05, n=737). Similar results were observed in analysis restricted to subjects with an HbA1c <= 5.7 (p_FWE_-corrected< .05, n= 712, please see Supplementary Results 2 and 3).

### 3.3. Cognitive performance and white matter microstructure

Within regions that showed a significant association of FA and HbA1c, we could show a negative correlation between global cognitive performance and FA in several clusters including the bilateral superior longitudinal fascicle as well as the bilateral corticospinal tract and the cerebellar peduncles (n= 1050, p_FWE_-corrected< .05). For more information please see Supplementary Results 4. Additional correlational analyses yielded similar associations between extracted FA values and cognitive subdomains with strongest effect sizes for the List Sorting Working Memory test (working memory) (n= 1050, r= .240, p< 0.001), the Picture Vocabulary test (vocabulary knowledge) (n= 1050, r= .185, p< 0.001) and the Penn Progressive Matrices test (fluid intelligence) (n= 1050, r= .140, p< 0.001). For an overview of the respective analyses of all cognitive subdomains, please see Table 1.

### 3.4. Analyses controlling for years of education

Total years of education were negatively correlated with HbA1c (n= 819, r= -.166, p< .001) and positively correlated with mean FA (n= 736, r= -.173, p= .027), as well as with global (n= 1180, r= .459, p< .001) cognition.

The effect size of the association between HbA1c and the global cognition score decreased when correcting for education years and thus failed to reach significance (df= 724, r= -.055, p= .139). However negative correlations between HbA1c and several cognitive subdomains including amongst others the List Sorting Working Memory Test (working memory) (df= 724, r= -0.089, p= .0.017), Picture Vocabulary Test (vocabulary knowledge) (df= 724, r= -0.076, p= 0.040) and the Penn Progressive Matrices (fluid intelligence) (df= 724, r= -0.080, p= 0.031) could still be observed after correction for education years (Supplementary Results 5). Likewise, the negative correlation between HbA1c and FA (df= 724, r= -.205, p< .001) as well as the positive correlation of FA and global cognition (df= 724, r= .200, p<.001) remained significant when correcting for education years. For detailed (education-years-corrected) correlational analyses with all cognitive measures, please see Supplementary Results 5.

**Table 1:** Correlation of HbA1c and FA with cognitive subscores

|  |  | HbA1C | FA |
| --- | --- | --- | --- |
| Picture Sequence Memory:<br>[non-verbal] episodic memory | Pearson's r | -0,047 | **,203\*\*** |
|  | p-value | 0,208 | **<0,001** |
|  | n | 730 | **1050** |
| Dimensional Change Card Sort Test:<br>executive function & cognitive flexibility | Pearson's r | -0,032 | **,112\*\*** |
|  | p-value | 0,395 | **0,000** |
|  | n | 730 | **1050** |
| Flanker Inhibitory Control and Attention Test:<br>executive function | Pearson's r | 0,063 | 0,058 |
|  | p-value | 0,087 | 0,061 |
|  | n | 730 | 1050 |
| Penn Progressive Matrices, total correct<br>responses:<br>fluid intelligence | Pearson's r | **-,118\*\*** | **,140\*\*** |
|  | p-value | **0,001** | **<0.001** |
|  | n | **728** | **1046** |
| Oral Reading Recognition Test:<br>reading decoding skills | Pearson's r | **-,086\*** | **,163\*\*** |
|  | p-value | **0,020** | **<0.001** |
|  | n | **730** | **1050** |
| Picture Vocabulary Test:<br>vocabulary knowledge | Pearson's r | **-,125\*\*** | **,185\*\*** |
|  | p-value | **0,001** | **<0.001** |
|  | n | **730** | **1050** |
| Pattern Comparison Processing Speed Test | Pearson's r | -0,052 | **,138\*\*** |
|  | p-value | 0,157 | **<0.001** |
|  | n | 730 | **1050** |
| Delay Discounting, 40k:<br>Self-regulation/ Impulsivity | Pearson's r | **-,100\*\*** | **,141\*\*** |
|  | p-value | **0,007** | **<0.001** |
|  | n | **729** | **1047** |
| Variable Short Penn Line Orientation, total correct items:<br>spatial orientation processing | Pearson's r | -0,046 | **,095\*\*** |
|  | p-value | 0,220 | **0,002** |
|  | n | 729 | **1047** |
| Short Penn Continuous Performance Test, specificity of right decisions:<br>Sustained attention | Pearson's r | -0,063 | **,123\*\*** |
|  | p-value | 0,091 | **<0.001** |
|  | n | 729 | **1047** |
| Penn Word Memory Test:<br>verbal episodic memory | Pearson's r | -0,064 | **,142\*\*** |
|  | p-value | 0,086 | **<0.001** |
|  | n | 729 | **1047** |
| List Sorting Working Memory Test | Pearson's r | **-,113\*\*** | **,152\*\*** |
|  | p-value | **0,002** | **<0.001** |
|  | n | **730** | **1050** |
| NIH Total Cognition Score | Pearson's r | **-,104\*\*** | **,240\*\*** |
|  | p-value | **0,005** | **<0.001** |
|  | n | **730** | **1050** |

HbA1c: Percentage of glycated Hemoglobin A1c; FA: Fractional anisotropy, mean value extracted from the FA-global cognition association results mask; Pearson's r= Pearson correlation coefficient; p-value= value determining statistical significance, \*\* p< 0.01; \* p< 0.05, values < .05 in bold font; n= number of subjects available for this distinct analysis; for more information on the cognitive subscores see Supplementary Methods 1

**Figure 1.**
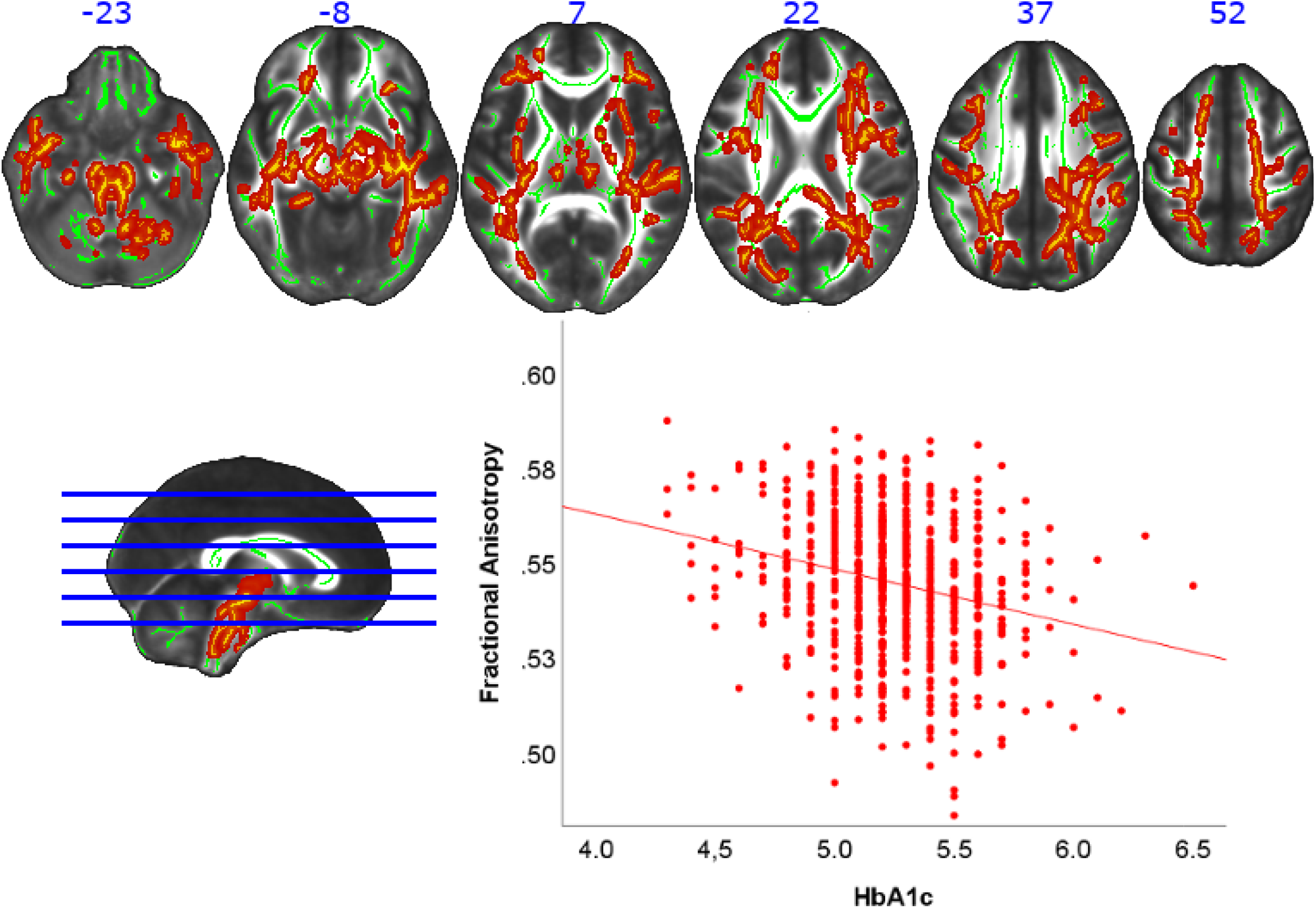
Negative association of HbA1c and Fractional Anisotropy. Top: Axial slices with corresponding y-axis values (MNI) are presented. Red-yellow areas represent voxels (using FSL’s, fill” command for better visualization), where a significant negative association between HbA1c and Fractional Anisotropy was detected (p_FWE_ < .05). Bottom left: Sagittal view with blue lines indicating axial slices on the top. Bottom right: Scatterplot showing the association of HbA1c and extracted mean FA values from all significant voxels of the corresponding TBSS analysis.

## 4 Discussion

With the present study we provide first evidence for a negative association between HbA1c and both cognitive performance and white matter microstructure in healthy, young adults. Our findings demonstrate that variation of HbA1c levels below the currently applied diagnostic cut-off scores for type II diabetes significantly affects cognition and brain structure. This finding underscores the idea of potential adverse effects of impaired glucose metabolism on cognition and brain structure that precede the actual onset of clinical manifest diabetes. According to the latter notion, preventive measures addressing nutrition habits and glucose metabolism should be reevaluated independently of the currently applied HBAlc cut-off scores for diabetes.

Although cross-sectional, results of the current analysis support a mechanistic concept in which subtle variations in HbA1c might impact white matter microstructure that in turn is highly likely to result in impaired cognitive function. This mechanistic concept heavily relies on evidence from multiple previous longitudinal and cross-sectional studies robustly demonstrating both associations between altered glucose metabolism, cognition and brain structure ^33^. There is substantial evidence from longitudinal studies in humans and animal studies pointing towards an effect of poor glycemic control on brain structure and cognitive function: In a longitudinal study (n=3469, mean age= 72.66 years) HbA1c levels were associated with greater episodic memory decline in diabetes patients, but not in healthy controls ^34^. Another longitudinal study in healthy elderly (>75 years of age) subjects showed a decrease of 1.37 points in the Mini Mental State Exam per unit increase in HbA1 at follow up (mean follow-up duration: 32 months) ^35^. In addition, evidence from animal models support the notion that hyperglycemia affects brain structure and cognition ^36,37^.

Regarding potential neurobiological pathways that could mediate the observed association between glucose metabolism and white matter integrity in healthy young subjects, biological processes beyond the classical concept of microangiopathy and macrovascular disease must be taken into consideration. While we cannot fully exclude that vascular damage might have contributed to the observed findings, the age distribution and lack of severe medical comorbidities in our sample warrant the discussion of further mechanisms that might help to explain the relationship between altered glucose metabolism and brain structural changes. One potential biological pathway that has frequently been purported as a potential mediator between metabolic deviation and brain structural damage involves aberrant immune response. The established connection between the immune system and metabolic disorders strongly supports this concept ^38^. Meta-analytic evidence confirms that inflammatory processes are involved in the pathophysiology of metabolic disorders including diabetes and obesity ^39,40^. Importantly, low grade peripheral inflammation has furthermore been associated with increased risk for diabetes ^39^. The latter finding might be of particular relevance for the interpretation of our findings that are derived from healthy non-diabetic participants. Both metabolic disorder and inflammation in turn have repeatedly been linked to brain structural alterations ^28,41-43^. More specifically, peripheral as well as CSF inflammatory markers have repeatedly been associated with impaired white matter microstructure in healthy subjects as well as in neuropsychiatric disorders ^44–46^. To sum up, in light of the current evidence, it appears highly suggestive that inflammatory processes might constitute one of several biological mechanisms potentially mediating the relationship between metabolic deviation and brain structural damage. Future studies should specifically test this hypothesis and clarify the precise underpinnings and causal directions using longitudinal designs. Further insights in this domain might be of great value for the future development of early interventions and preventive strategies.

In the affected withe matter tracts, we could demonstrate a negative association between FA and global cognitive performance. Although our results were widespread and included both superior longitudinal fascicles as well as the bilateral corticospinal tract, the largest cluster was found in the cerebellar peduncles. Interestingly, this is in line with previous evidence supporting a crucial role of the cerebellum in cognitive performance (for a review see ^47^) and suggests disturbed cortical-cerebellar neuronal communication as a possible mechanism for the adverse effect of HbA1c on cognition.

Another important finding of the present study is that associations between HbA1c and brain structure as well as between HbA1c and cognitive performance withstands correction for important covariates that are likely to covary with either HbA1c, cognition or brain structure. More precisely, our results appeared to be independent of age and BMI, two important sources of potential bias regarding the studied independent and dependent variables ^28,48^. Furthermore, as could be expected the effect size of the observed associations decreased in analyses controlling for education. This is not surprising considering the strong impact of education on both cognition and brain structure ^49^, as well as the suspected association between education/ socioeconomic factors and diet habits ^50^. However, even after controlling for the detrimental impact of this important socioeconomic variable, still significant associations between HbA1c and FA as well as between FA and cognition scores could be detected. These results confirm the observed association between HbA1c and brain structure and cognition and speak against possibility that our observations were mainly the result of spurious correlations or biased by undiscovered socioeconomic factors.

To further elucidate which subdomains of global cognitive performance drive the main findings of HbA1c-cognition and FA-cognition association, we performed exploratory correlational analyses with scores from all available cognitive measures available in HCP dataset. Here we found that several distinct subscores of cognitive performance were associated both with HbA1c as well as FA values (Table 1). A variety of cognitive tests that are considered to reflect both crystal cognition (Oral Reading Recognition Test, Picture Vocabulary Test) as well as fluid cognition (Penn Progressive Matrices, List Sorting Working Memory Test) are all significantly negatively associated with HbA1c and FA scores. This supports the concept of overall and global cognitive performance deficits (in contrast to a single distinct subscore driving the significant global cognition associations).

Besides these considerations, the results of our study hold several further important implications for future research. The core finding of the present study is that variation of HbA1c below the currently applied diagnostic criteria already associates with impaired cognitive function and brain structure. Most importantly, only n= 25 subjects exceeded HbA1c concentrations of 5.7%, and analyses excluding these subjects yielded highly similar results. Regarding this observation, the role of diagnostic cut off scores is challenged - a notion that is supported by the American Diabetes Association that proposed even stricter HbA1c cutoffs for people at risk than the WHO ^2,3^. In line with these reports, our results speak for a continuous rather than a categorical application of metabolic markers. In this regard, one should also take into consideration that the present study applied measures of cognition and white matter microstructure as assessed using MRI, measures that are highly likely the result of biological processes with an onset years before assessment.

Major strengths of the present study comprise the large sample size as well as the availability of both structural MRI and cognitive data allowing to infer on different levels biological abstraction. Furthermore, the available data allowed us to differentiate between a variety of cognitive measures.

A major limitation of the present study lies in the cross-sectional design that prevents us from assessing causal relationships. In this regard, it appears important to emphasize that the results of our study should be interpreted with caution especially when considering potential mechanistic relationships. Future longitudinal studies are needed to clarify the temporal precedence and potential causal relationships between glucose metabolism and altered brain structure and function. Furthermore, due to the lack of experimental conditions, our results risk to be biased by potential spurious correlations. However, it appears important to consider that in the present work, results of all main analyses withstood correction for common nuisance variables that were a priori likely to bias the observed associations including age, sex, BMI and socioeconomic factors like education.

To conclude the present study demonstrates an association between HbA1c concentrations below diagnostic cut off points for type II diabetes and white matter as well as cognitive performance in healthy, young adults. Our findings thus suggest a detrimental impact of subtle alterations in glucose metabolism on impaired brain structure and cognitive function. These results highlight the importance to further elucidate the potential mechanisms leading to neural damage in impaired glucose metabolism as well as to reconsider the current concept of HbA1c in diagnostics of diabetes. Our findings heavily support the notion of a continuous adverse impact of altered glucose metabolism on brain physiology that is likely to take its origin far below currently applied diagnostic cut-off scores. Future research and preventive measures should take these findings into consideration.

## ACKNOWLEDGMENTS

This work was funded by the German Research Foundation (DFG, grant FOR2107 DA1151/5-1 and DA1151/5-2 to UD; SFB-TRR58, Projects C09 and Z02 to UD) and the Interdisciplinary Center for Clinical Research (IZKF) of the medical faculty of Münster (grant Dan3/012/17 to UD).

Data were provided [in part] by the Human Connectome Project, WU-Minn Consortium (Principal Investigators: David Van Essen and Kamil Ugurbil; 1U54MH091657) funded by the 16 NIH Institutes and Centers that support the NIH Blueprint for Neuroscience Research; and by the McDonnell Center for Systems Neuroscience at Washington University.

## Author contributions

JR and GK made substantial contributions to the conception, design of the work, and drafted the work. SM, KF, DG, JG made substantial contributions to the analysis of this work and the interpretation of data. VA BB UD have substantially revised the work. NO made substantial contributions to the conception, design of the work, and drafted the work.

All authors approved the submitted version and agreed both to be personally accountable for the author’s own contributions and that questions related to the accuracy or integrity of any part of the work are appropriately investigated.

---

**Supplementary Methods 1:**
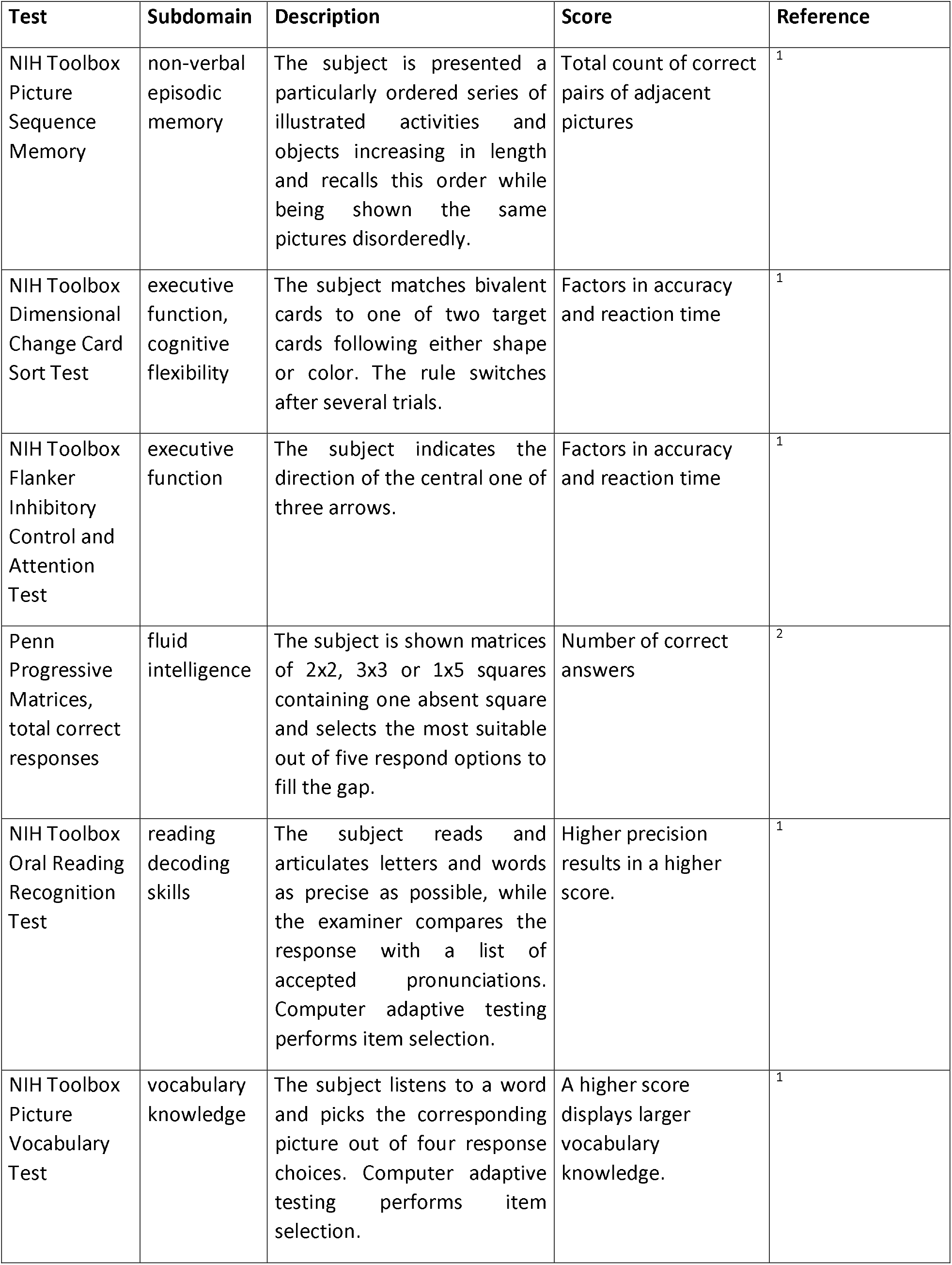
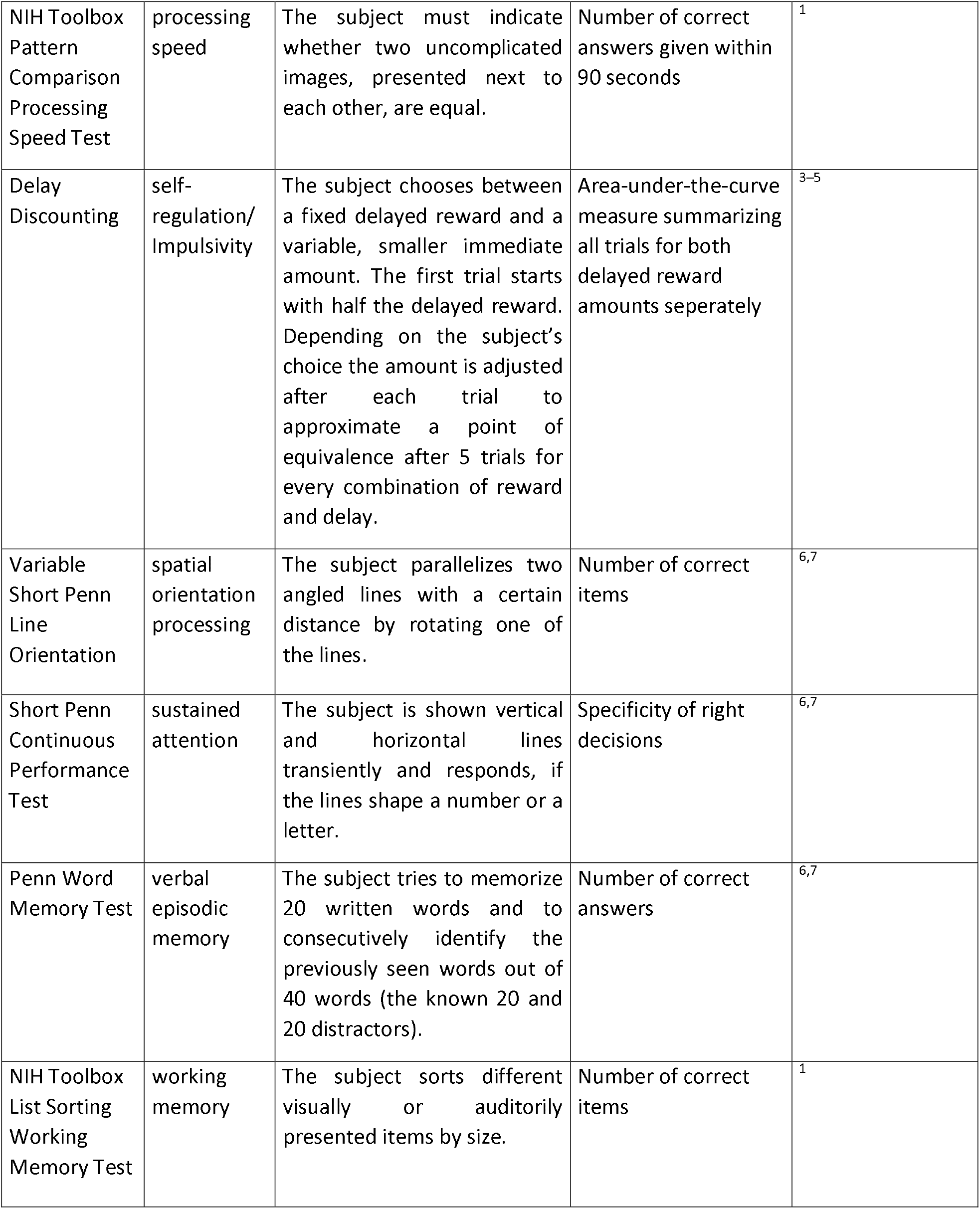
Neurocoenitive Tests in the HCP sample

| **Test** | **Subdomain** | **Description** | **Score** | **Reference** |
| --- | --- | --- | --- | --- |
| NIH Toolbox Picture Sequence Memory | non-verbal episodic memory | The subject is presented a particularly ordered series of illustrated activities and objects increasing in length and recalls this order while being shown the same pictures disorderedly. | Total count of correct pairs of adjacent pictures | <sup>1</sup> |
| NIH Toolbox Dimensional Change Card Sort Test | executive function, cognitive flexibility | The subject matches bivalent cards to one of two target cards following either shape or color. The rule switches after several trials. | Factors in accuracy and reaction time | <sup>1</sup> |
| NIH Toolbox Flanker Inhibitory Control and Attention Test | executive function | The subject indicates the direction of the central one of three arrows. | Factors in accuracy and reaction time | <sup>1</sup> |
| Penn Progressive Matrices, total correct responses | fluid intelligence | The subject is shown matrices of 2x2, 3x3 or 1x5 squares containing one absent square and selects the most suitable out of five respond options to fill the gap. | Number of correct answers | <sup>2</sup> |
| NIH Toolbox Oral Reading Recognition Test | reading decoding skills | The subject reads and articulates letters and words as precise as possible, while the examiner compares the response with a list of accepted pronunciations. Computer adaptive testing performs item selection. | Higher precision results in a higher score. | <sup>1</sup> |
| NIH Toolbox Picture Vocabulary Test | vocabulary knowledge | The subject listens to a word and picks the corresponding picture out of four response choices. Computer adaptive testing performs item selection. | A higher score displays larger vocabulary knowledge. | <sup>1</sup> |
| NIH Toolbox Pattern Comparison Processing Speed Test | processing speed | The subject must indicate whether two uncomplicated images, presented next to each other, are equal. | Number of correct answers given within 90 seconds | <sup>1</sup> |
| Delay Discounting | self-regulation/ Impulsivity | The subject chooses between a fixed delayed reward and a variable, smaller immediate amount. The first trial starts with half the delayed reward. Depending on the subject's choice the amount is adjusted after each trial to approximate a point of equivalence after 5 trials for every combination of reward and delay. | Area-under-the-curve measure summarizing all trials for both delayed reward amounts separately | <sup>3-5</sup> |
| Variable Short Penn Line Orientation | spatial orientation processing | The subject parallelizes two angled lines with a certain distance by rotating one of the lines. | Number of correct items | <sup>6,7</sup> |
| Short Penn Continuous Performance Test | sustained attention | The subject is shown vertical and horizontal lines transiently and responds, if the lines shape a number or a letter. | Specificity of right decisions | <sup>6,7</sup> |
| Penn Word Memory Test | verbal episodic memory | The subject tries to memorize 20 written words and to consecutively identify the previously seen words out of 40 words (the known 20 and 20 distractors). | Number of correct answers | <sup>6,7</sup> |
| NIH Toolbox List Sorting Working Memory Test | working memory | The subject sorts different visually or auditorily presented items by size. | Number of correct items | <sup>1</sup> |

**Supplementary Results 1:** Whole Sample: Negative correlation of Fractional Anisotropy with HbA1c (n=737)

| Voxels | $p_{FWE}$ | MNI X (mm) | MNI Y (mm) | MNI Z (mm) |
| --- | --- | --- | --- | --- |
| 25241 | 0.001 | 5 | -28 | -19 |
| 10487 | 0.019 | -35 | -47 | 29 |
| 678 | 0.047 | -17 | -73 | 14 |
| 283 | 0.047 | -21 | 43 | 15 |

## Probabilities of affected tracts in percent

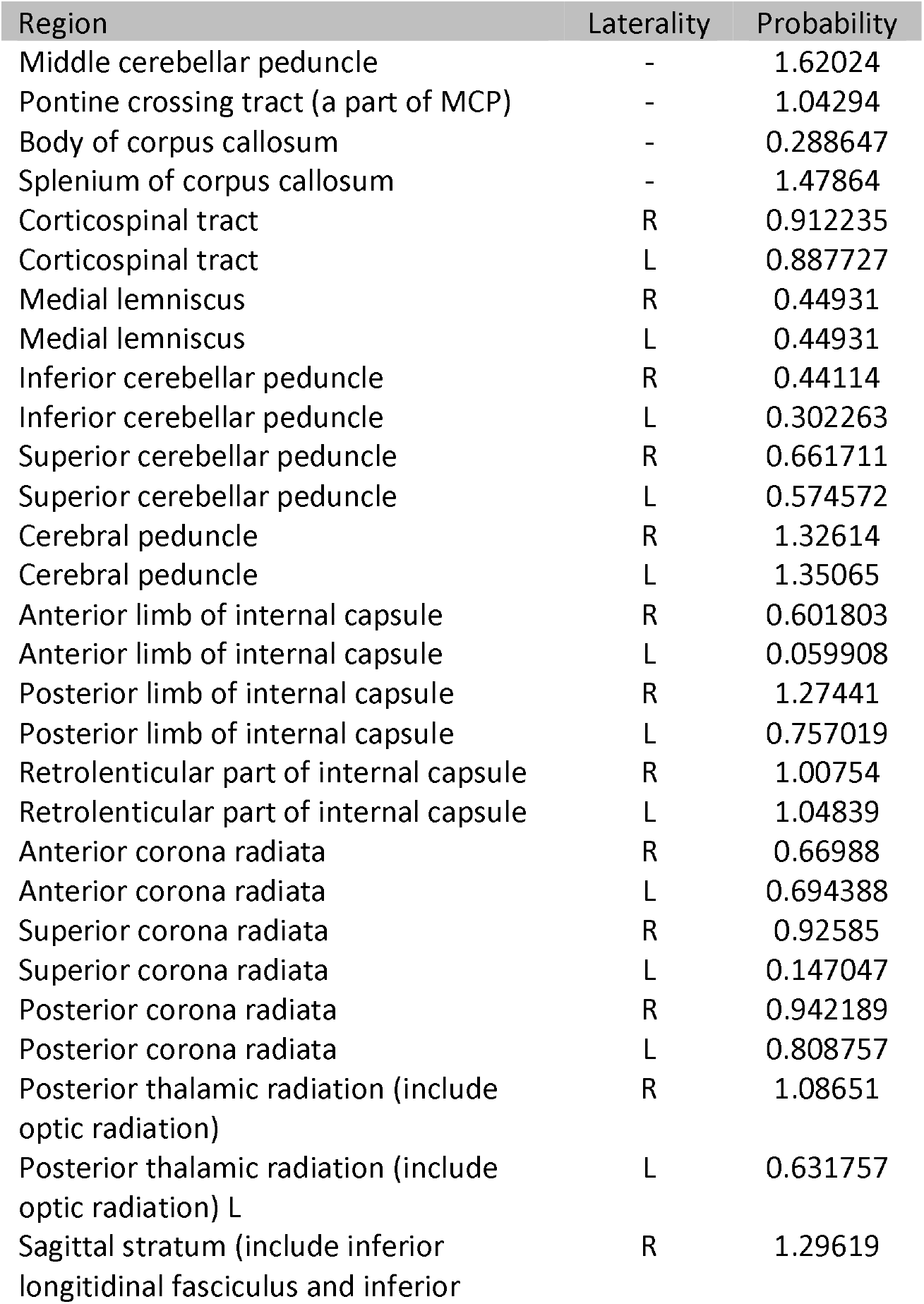
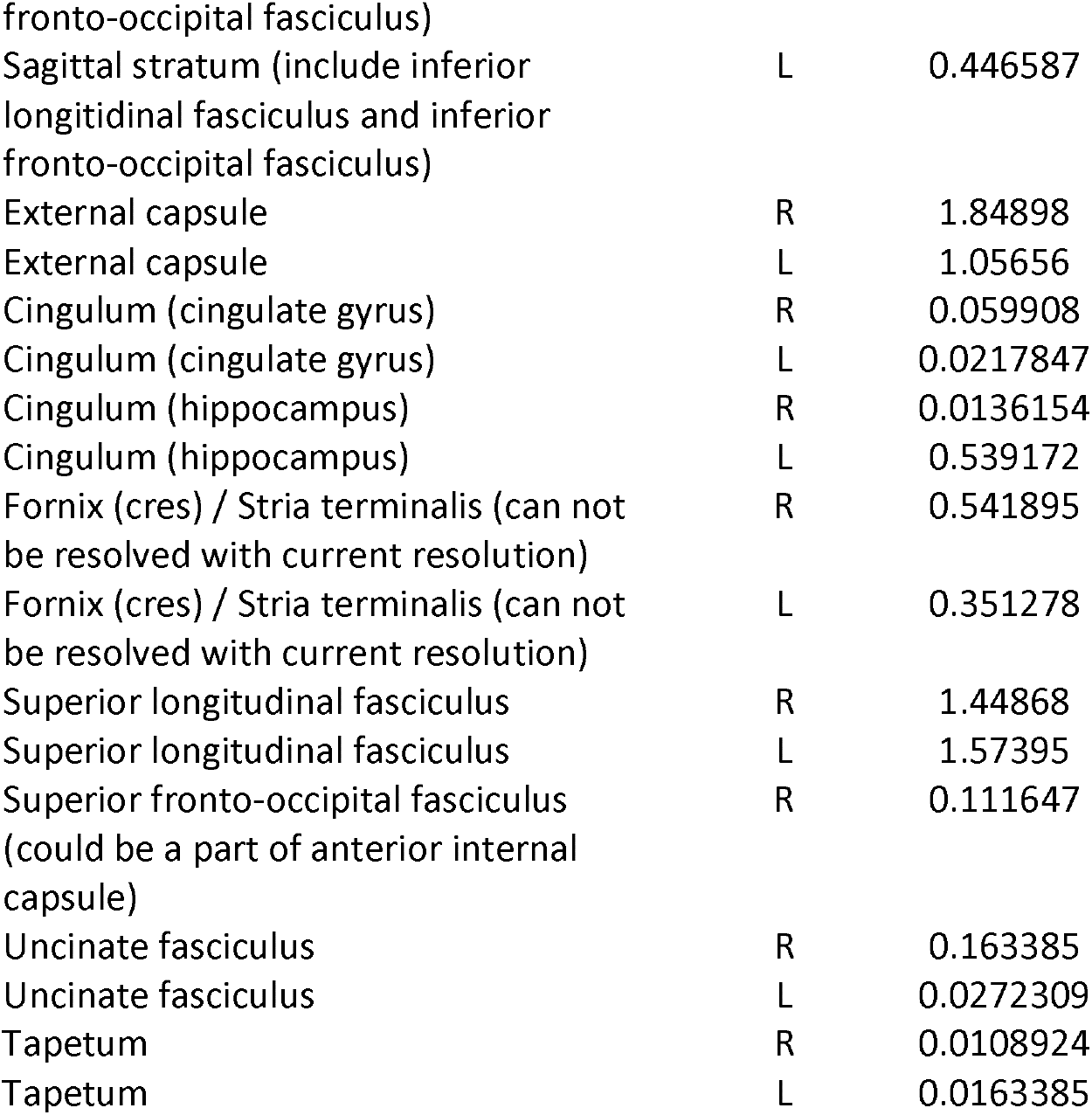

### Positive correlation

No significant results

On the top dimensions of clusters (number of voxels) and localization of signal peaks (MNI coordinates) are given for regions showing maximal differences of tract-based spatial statistics values (signal peak). Below are the white matter tracts in the cluster based on the JHU ICBM-DTI-81 White-Matter Labels (as implemented in FSL).

Probabilities of affected tracts: It gives the (average) probability of all significant voxels being a member of the different labelled regions within the atlas (JHU ICBM-DTI-81 White-Matter), calculated with the FSL tool “atlasquery”

**Supplementary Results 2:** Sample with HbA1c =< 5.7: Negative correlation of Fractional Anisotropy with HbA1c (n=712)

| Voxels | $p_{FWE}$ | MNI X (mm) | MNI Y (mm) | MNI Z (mm) |
| --- | --- | --- | --- | --- |
| 45315 | .001 | -3 | -28 | -23 |
| 85 | 0.049 | 18 | -79 | 18 |

## Probabilities of affected tracts in percent

**Probabilities of affected tracts in percent:**
| Region | Laterality | Probability |
| --- | --- | --- |
| Middle cerebellar peduncle | - | 1.49119 |
| Pontine crossing tract (a part of MCP) | - | 0.80837 |
| Genu of corpus callosum | - | 1.23568 |
| Body of corpus callosum | - | 3.0793 |
| Splenium of corpus callosum | - | 2.18722 |
| Corticospinal tract R | R | 0.707048 |
| Corticospinal tract L | L | 0.744493 |
| Medial lemniscus R | R | 0.35022 |
| Medial lemniscus L | L | 0.334802 |
| Inferior cerebellar peduncle R | R | 0.339207 |
| Inferior cerebellar peduncle L | L | 0.284141 |
| Superior cerebellar peduncle R | R | 0.555066 |
| Superior cerebellar peduncle L | L | 0.486784 |
| Cerebral peduncle R | R | 1.06608 |
| Cerebral peduncle L | L | 1.14758 |
| Anterior limb of internal capsule R | R | 0.814978 |
| Anterior limb of internal capsule L | L | 0.0506608 |
| Posterior limb of internal capsule R | R | 1.09912 |
| Posterior limb of internal capsule L | L | 0.632159 |
| Retrolenticular part of internal capsule R | R | 0.914097 |
| Retrolenticular part of internal capsule L | L | 1.00661 |
| Anterior corona radiata R | R | 1.17181 |
| Anterior corona radiata L | L | 1.10573 |
| Superior corona radiata R | R | 0.759912 |
| Superior corona radiata L | L | 0.396476 |
| Posterior corona radiata R | R | 0.995595 |
| Posterior corona radiata L | L | 0.685022 |
| Posterior thalamic radiation (include optic radiation) R | R | 0.788546 |
| Posterior thalamic radiation (include optic radiation) L | L | 0.731278 |
| Sagittal stratum (include inferior longitudinal fasciculus and inferior fronto-occipital fasciculus) R | R | 0.997797 |
| Sagittal stratum (include inferior longitudinal fasciculus and inferior fronto-occipital fasciculus) L | L | 0.319383 |
| External capsule R | R | 1.61454 |
| External capsule L | L | 1.03304 |
| Cingulum (cingulate gyrus) R | R | 0.259912 |
| Cingulum (cingulate gyrus) L | L | 0.0176211 |
| Cingulum (hippocampus) R | R | 0.0110132 |
| Cingulum (hippocampus) L | L | 0.462555 |
| Fornix (cres) / Stria terminalis (can not be resolved with current resolution) R | R | 0.453745 |
| Fornix (cres) / Stria terminalis (can not be resolved with current resolution) L | L | 0.559471 |
| Superior longitudinal fasciculus R | R | 1.49559 |
| Superior longitudinal fasciculus L | L | 1.44714 |
| Superior fronto-occipital fasciculus (could be a part of anterior internal capsule) R | R | 0.105727 |
| Uncinate fasciculus R | R | 0.132159 |
| Uncinate fasciculus L | L | 0.0220264 |
| Tapetum R | R | 0.0418502 |
| Tapetum L | L | 0.0154185 |

### Positive correlation

No significant results.

**Supplementary Results 3:** Whole Sample: Negative correlation of Fractional Anisotropy with HbA1c corrected for body mass index (n=737)

**corrected for body mass index (n=737)**
| Voxels | $p_{FWE}$ | MNI X (mm) | MNI Y (mm) | MNI Z (mm) |
| --- | --- | --- | --- | --- |
| 3579 | .011 | 7 | -22 | -15 |
| 1084 | .036 | 40 | -20 | -8 |
| 873 | .046 | 26 | -39 | 30 |
| 218 | .046 | -32 | -50 | 26 |
| 156 | .046 | 36 | 39 | -3 |
| 67 | .048 | 27 | -65 | 28 |
| 62 | .049 | 16 | -16 | 58 |
| 59 | .049 | -18 | -66 | -34 |

## Probabilities of affected tracts in percent

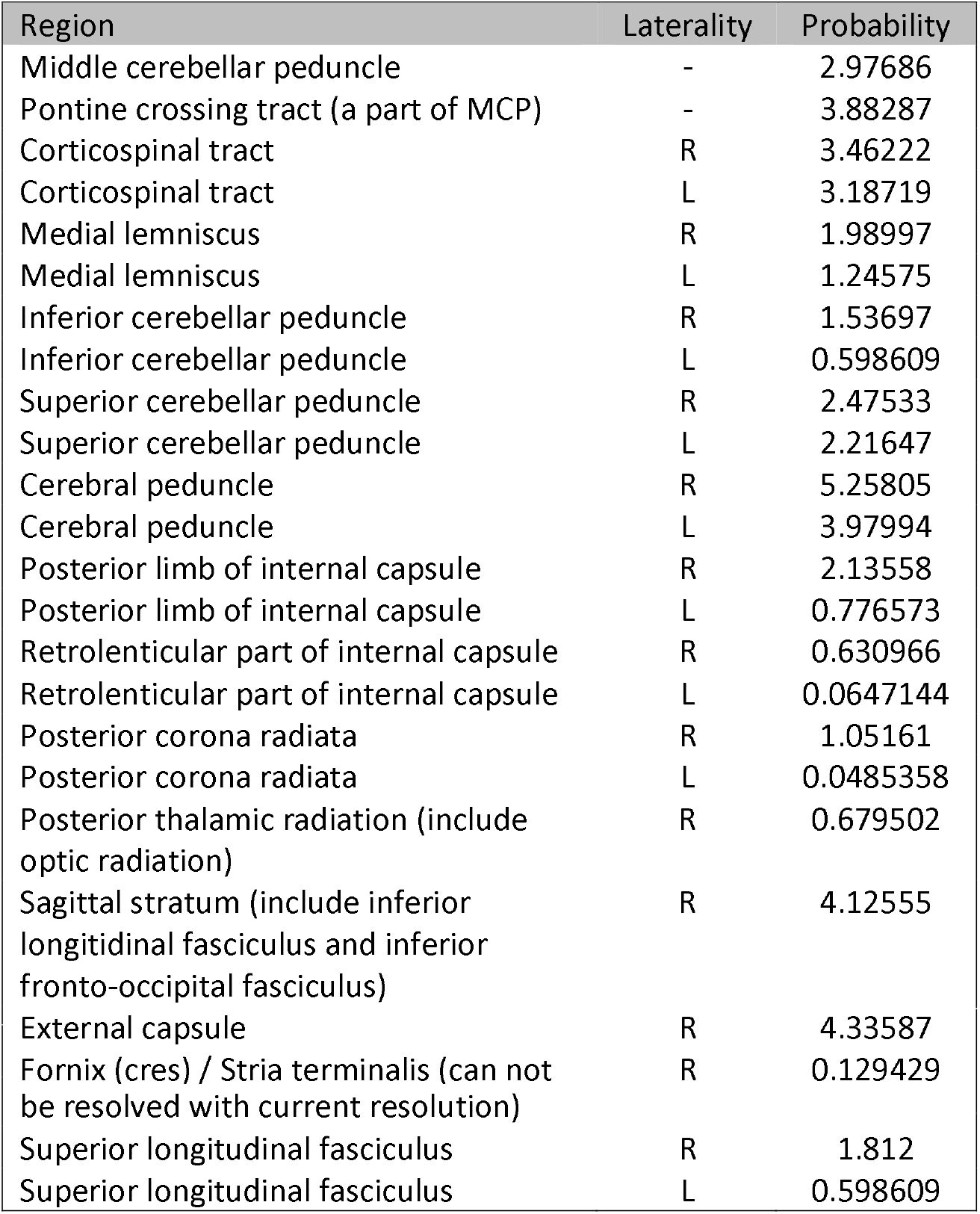

### Positive correlation

No significant results.

**Results 4:** Positive Correlation of Fractional Anisotropy with global cognitive performance (n= 1050)

| Voxels | $p_{FWE}$ | MNI X (mm) | MNI Y (mm) | MNI Z (mm) |
| --- | --- | --- | --- | --- |
| 2014 | < .001 | 7 | -41 | -30 |
| 370 | .005 | -36 | -44 | 30 |
| 226 | .012 | 30 | -36 | 37 |
| 188 | .021 | 33 | -49 | 34 |
| 154 | .018 | -36 | -4 | -24 |
| 95 | 0.002 | -35 | -20 | 34 |
| 77 | 0.014 | -20 | -66 | -37 |
| 77 | 0.028 | 34 | -13 | -4 |
| 54 | 0.042 | 30 | -17 | -9 |

## Probabilities of affected tracts in percent

**Probabilities of affected tracts in percent:**
| Region | Laterality | Probability |
| --- | --- | --- |
| Middle cerebellar peduncle | - | 4.44243 |
| Pontine crossing tract (a part of MCP) | - | 0.453309 |
| Corticospinal tract | R | 4.68419 |
| Corticospinal tract | L | 4.9864 |
| Medial lemniscus | R | 2.96162 |
| Medial lemniscus | L | 2.35721 |
| Inferior cerebellar peduncle | R | 1.05772 |
| Inferior cerebellar peduncle | L | 0.604412 |
| Superior cerebellar peduncle | R | 6.22545 |
| Superior cerebellar peduncle | L | 3.98912 |
| Cerebral peduncle | R | 3.68691 |
| Cerebral peduncle | L | 4.50287 |
| Posterior limb of internal capsule | L | 0.0906618 |
| Posterior corona radiata | R | 0.0302206 |
| Sagittal stratum (include inferior longitudinal fasciculus and inferior fronto-occipital fasciculus) | R | 0.120882 |
| Sagittal stratum (include inferior longitudinal fasciculus and inferior fronto-occipital fasciculus) | L | 0.0302206 |
| External capsule | R | 2.32699 |
| Fornix (cres) / Stria terminalis (can not be resolved with current resolution) | R | 0.815956 |
| Fornix (cres) / Stria terminalis (can not be resolved with current resolution) | L | 0.211544 |
| Superior longitudinal fasciculus | R | 4.56331 |
| Superior longitudinal fasciculus | L | 5.07706 |

### Negative correlation

No significant results.

Analyses were performed in all regions that showed a negative association of HbA1c and fractional anisotropy (Supplementary Results 1).

**Supplementary Results 5:** Correlation of HbA1c and FA with cognitive subscores corrected for total education years

|  |  | HbA1C | FA |
| --- | --- | --- | --- |
| Picture Sequence Memory:<br>[non-verbal] episodic memory | Pearson's r | -0,023 | 0,188 |
|  | p-value | 0,533 | 0,000 |
|  | df | 724 | 724 |
| Dimensional Change Card Sort Test:<br>executive function & cognitive flexibility | Pearson's r | -0,014 | 0,087 |
|  | p-value | 0,712 | 0,020 |
|  | df | 724 | 724 |
| Flanker Inhibitory Control and Attention Test:<br>executive function | Pearson's r | 0,075 | 0,027 |
|  | p-value | 0,044 | 0,466 |
|  | df | 724 | 724 |
| Penn Progressive Matrices, total correct<br>responses:<br>fluid intelligence | Pearson's r | -0,080 | 0,098 |
|  | p-value | 0,031 | 0,008 |
|  | df | 724 | 724 |
| Oral Reading Recognition Test:<br>reading decoding skills | Pearson's r | -0,034 | 0,111 |
|  | p-value | 0,364 | 0,003 |
|  | df | 724 | 724 |
| Picture Vocabulary Test:<br>vocabulary knowledge | Pearson's r | -0,076 | 0,114 |
|  | p-value | 0,040 | 0,002 |
|  | df | 724 | 724 |
| Pattern Comparison Processing Speed Test | Pearson's r | -0,035 | 0,125 |
|  | p-value | 0,342 | 0,001 |
|  | df | 724 | 724 |
| Delay Discounting, 40k:<br>Self-regulation/ Impulsivity | Pearson's r | -0,073 | 0,106 |
|  | p-value | 0,048 | 0,004 |
|  | df | 724 | 724 |
| Variable Short Penn Line Orientation, total correct items:<br>spatial orientation processing | Pearson's r | -0,023 | 0,049 |
|  | p-value | 0,536 | 0,192 |
|  | df | 724 | 724 |
| Short Penn Continuous Performance Test, specificity of right decisions:<br>Sustained attention | Pearson's r | -0,042 | 0,094 |
|  | p-value | 0,260 | 0,011 |
|  | df | 724 | 724 |
| Penn Word Memory Test:<br>verbal episodic memory | Pearson's r | -0,039 | 0,153 |
|  | p-value | 0,288 | 0,000 |
|  | df | 724 | 724 |
| List Sorting Working Memory Test | Pearson's r | -0,089 | 0,154 |
|  | p-value | 0,017 | 0,000 |
|  | df | 724 | 724 |
| NIH Total Cognition Score | Pearson's r | -0,055 | 0,200** |
|  | p-value | 0,139 | 0,000 |
|  | df | 724 | 724 |

HbA1c: Percentage of glycated Hemboglobin A1c
FA: Fractional anisotropy, mean value extracted from the FA-global cognition association results mask
\*\* p< 0.01
\* p< 0.05
For more information on the cognitive subscores see Supplementary Methods 1.

